# Likelihood-based docking of models into cryo-EM maps

**DOI:** 10.1101/2022.12.20.521188

**Authors:** Claudia Millán, Airlie J. McCoy, Thomas C. Terwilliger, Randy J. Read

## Abstract

Optimized docking of models into cryo-EM maps requires exploiting an understanding of the signal expected in the data to minimize the calculation time while maintaining sufficient signal. The likelihood-based rotation function used in crystallography can be employed to establish plausible orientations in a docking search. A phased likelihood translation function yields scores for the placement and rigid-body refinement of oriented models. Optimised strategies for choices of the resolution of data from the cryo-EM maps to use in the calculations and the size of search volumes are based on expected log-likelihood-gain scores, computed in advance of the search calculation. Tests demonstrate that the new procedure is fast, robust and effective at placing models into even challenging cryo-EM maps.

**Synopsis:** Exploiting analogies to crystallographic molecular replacement, a strategy for docking into cryo-EM maps is informed by calculations of expected log-likelihood-gain scores.

## 1. Introduction

Advances in cryo-EM hardware and software are improving the resolution and quality of cryo-EM maps, often yielding maps that allow model-building from scratch. Nevertheless, for various sample-specific and technical reasons, a substantial proportion of cryo-EM maps from single-particle reconstructions and a larger proportion of maps from sub-tomogram averaging lack the necessary resolution and quality for *ab initio* model building. In this situation, the density may be interpreted by docking one or more pre-existing experimental or predicted atomic models. We explore here the development of a new likelihood-based docking tool, *em_placement*, to fill this need. The accompanying paper (Read *et al*., 2023) builds the theoretical background for the required likelihood targets and the statistical calculations underlying automation features of this software.

A large number of tools have been developed to carry out manual or automated docking. The automated tools include DockEM (Roseman, 2000; Titarenko & Roseman, 2021), Situs (Kovacs & Wriggers, 2002; Wriggers, 2012), Powerfit (Zundert *et al*., 2015), OffGridFit (Hoffmann *et al*., 2017), phenix.dock_in_map (Liebschner *et al*., 2019), and MrBUMP (Simpkin *et al*., 2021). DockEM, Powerfit and phenix.dock_in_map all carry out an exhaustive exploration of orientations. Situs and OffGridFit both use 6-dimensional FFT-based algorithms for an exhaustive 6D search. Among these, MrBUMP is unique in carrying out the translation search first with the spherically-averaged phased translation function, followed by an orientation search centered on the point found in the translation search.

We have not attempted to carry out head-to-head comparisons of our software with existing tools for two reasons. First, the half-maps needed for our approach are not generally available for the published test cases for existing tools. Second, we are not experts in the use of the other tools and would therefore not be able to show them to their best advantage.

## 2. Expected LLG-based search strategy

Docking problems can differ dramatically in their difficulty, from trivial cases where distinctive features of the search model could be spotted by eye in an excellent map, to extremely challenging cases where there is barely enough signal to recognise that a docked model agrees with a very noisy map. Great gains can be made in the efficiency and effectiveness of docking calculations by adopting a case-dependent strategy that is informed by considering the value of the log-likelihood-gain (LLG) score that would be expected for a correct solution given the quality of the data and the model, which we term the expected LLG or eLLG. The accompanying paper (Read *et al*., 2023) provides the details of how these are defined and computed.

In crystallographic molecular replacement (MR), we have found that searches yielding an LLG value of 60 or greater after a combined rotation/translation search are almost always correct (McCoy *et al*., 2017; Oeffner *et al*., 2018). In cryo-EM, after correcting for oversampling, our experiences in the tests below suggest that a similar threshold applies for identifying correct, or at least non-random, solutions. Given uncertainties about the sizes of coordinate errors prior to structure solution, trials of different choices in a database of MR problems showed that it is more efficient, overall, to choose strategy parameters expected to give a higher LLG score than 60, with 225 being a choice that works well to balance an increased initial search cost with a lower chance of having to rerun an unsuccessful search with modified parameters.

The pivotal decisions in the docking search strategy are determined by the rotation search, because it gives the lowest signal-to-noise; if this search is expected to succeed (or at least to give sufficient signal that a chosen subset of orientations is likely to include the correct orientation), then the subsequent translation search will be almost certain to succeed.

For this reason, the strategy decisions discussed below are primarily driven by considerations of the LLG signal expected in the rotation search, the *eLLG*_*rot*_. The *eLLG*_*rot*_ depends on the resolution of the map and the fraction of the ordered volume in that map explained by the model. For a model covering only a small part of the ordered volume in a low-resolution map, the *eLLG*_*rot*_ signal can be very low. Fortunately, cryo-EM differs from crystallography in that the phase information allows a search to be confined to only a part of the full map, which reduces noise in the rotation search and increases *eLLG*_*rot*_.

Given uncertainties in the eLLG calculations, particularly in estimating the quality of the search model in advance, searches that aim for the minimum required LLG have a significant chance of failure, and it is safer to be somewhat more conservative so that fewer searches need to be repeated. For the rotation search, an LLG score of 30 or more is expected to correspond to a correct solution, as this is equivalent to the search score required for confidence in a crystallographic MR search in space group P1 (McCoy *et al*., 2017). As a more conservative estimate, the initial target *eLLG*_*rot*_ is set to 60.

### 2.1 Searching over the whole map with one rotation search

A major decision in the search strategy is whether a rotation search over the whole map is likely to succeed. For good maps and good models that comprise a sufficient fraction of the total structure, a strong signal will be expected in the rotation search. Searches can then be carried out over the whole map, but the efficiency is optimised, by default in *em_placement*, by limiting the resolution to what is required to achieve *eLLG*_*rot*_ = 60. This is a value that will usually yield an unambiguous and precise orientation. In principle, an even lower resolution limit for Fourier terms could be used in the translation search, but in practice the translation search is computationally very efficient, and only a few translation searches will be required if there is good signal in the rotation search. When it is not possible to achieve the ideal value of *eLLG*_*rot*_ = 60, lower levels of signal are still useful. Even if *eLLG*_*rot*_ = 7.5, the correct orientation is likely to be found in an orientation list of modest size, so carrying out a rotation search followed by a translation search over the entire map is only abandoned if *eLLG*_*rot*_ < 7.5.

### 2.2 Searching over sub-volumes

If it is not judged possible to search successfully for rotations using the full map, a decision is made whether it will be possible, instead, to find a solution by searching over sub-volumes. The target sub-volume is set according to the inverse relationship (Read *et al*., 2023) between the size of the sub-volume and the *eLLG*_*rot*_ that would be achieved by searching in that sub-volume (if it contained the object being sought). As a simple example, if a value of 3.75 were found for *eLLG*_*rot*_ when computed over the whole map, the *eLLG*_*rot*_ for a map containing one-half of the total volume would be 7.5, which is the default for the minimal acceptable value. This calculation depends on the assumption that one of the sub-volumes will contain the entire object being sought, so there is a lower limit to the smallest relevant sub-volume. It is also implicitly assumed that the map quality in local regions is not much worse than the overall average map quality. This can lead to failures when the component being sought corresponds to a poor part of the map. Note that there is a practical limit to how small a sub-volume can be; the number of overlapping sub-volumes required to ensure that at least one of them contains the full volume of the model grows dramatically once the search volume is less than about 1.15 times the volume of the sphere enclosing the model. When the required search volume would be unfeasibly small, the brute-force search discussed below is invoked.

When a suitable size has been defined for the sub-volumes (*i*.*e*. a size expected to achieve the minimal *eLLG*_*rot*_ of 7.5), target sub-volumes for docking searches are constructed as follows. First, a hexagonal close-packed grid is defined, such that spheres with the target volume that are centered on the grid points will overlap sufficiently that at least one of the spheres is guaranteed to cover the volume containing the target object. Second, any spheres that lack sufficient ordered volume (defined as regions of the map with high local variance) to contain the search object are discarded. Following this, the spheres of density are analysed to evaluate signal and noise (to calibrate the likelihood targets), using the program *prepare_map_for_docking* described in the accompanying paper (Read *et al*., 2023), and then rotation and translation searches are carried out, followed by rigid-body refinement. To avoid Fourier artefacts from sharp boundaries in the map, the target sphere is cut out inside a cube large enough to allow a smooth masking of the density to the edges.

### 2.3 Brute-force six-dimensional search

If the rotation search cannot be carried out with sufficient signal even with sub-volumes, then the final fall-back in the search algorithm is to carry out a brute-force six-dimensional search. To make this search as efficient as possible, data are used only to the resolution required to obtain a value of *eLLG*_*tra*_ sufficient to yield a clear solution for the correct combined rotation and translation. Based on experiences with crystallographic MR, searches given an LLG of 60 should almost always be correct, but to be safe the target for *eLLG*_*tra*_ is set to 225, a value that has also been adopted for crystallographic MR in *Phaser* to give a good compromise between efficiency and the danger of missing the solution. Using the lowest resolution possible improves efficiency by allowing orientations and translations to be sampled more coarsely, and by reducing the number of Fourier terms over which the likelihood scores must be calculated. Even so, it is not uncommon for such a brute-force search to take hours to run.

### 2.4 Focused docking

The final step in the docking strategy is to evaluate all potential docking solutions in a common framework. Docking poses that have an LLG score within some tolerance of the top score are retained as potential solutions. The tolerance is chosen by approximating the standard deviation of the LLG score as the square root of that score (McCoy *et al*., 2017) and allowing potential solutions to deviate by as much as 7 times that standard deviation. For each potential solution, the size of the sphere of density required to accommodate the entire search model is evaluated, a sphere of density of that size is cut out, the analysis of signal and noise is performed, and then a rigid-body refinement is carried out to obtain an LLG score, a final model placement and a map correlation with the processed density sphere. After the focused docking calculation, the list is pruned again based on the new top LLG score. By default, a maximum of 5 potential solutions are retained.

Two types of map coefficients (equations 18 and 19 from the accompanying paper, (Read *et al*., 2023)) for the processed density sphere have been evaluated in the set of tests described. The first type (**F**_*map*_ = *D*_*obs*_ **E**_*mean*_) should give a map that minimises the error from the true sharpened map because it represents the expected value of the Fourier coefficient for such a map. The second type 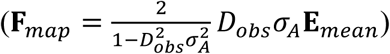 includes an additional weighting term from the likelihood target and therefore gives a map for which the correlation to a sharpened map computed from the docked model should be roughly proportional to the likelihood score for that model. To compute the second map, a choice has to be made for the value of *σ*_*A*_. The current default is to assign a value of 0.9, which would correspond to a model that accounts for about 80% of the scattering in the volume under consideration but has no other errors. The choice for *σ*_*A*_ could potentially be improved by considering deficiencies in the ability of atomic models to account for the bulk solvent region. The second choice for map coefficients yielded higher map correlations than the first in the test calculations reported below.

Qualitatively, the blurring that comes from giving higher weight to well-determined (typically lower-resolution) Fourier terms seems to give more readily interpretable maps, in line with the map correlation values. The second choice, therefore, is the default and was used for the map correlation calculations reported below.

## 3. Methods

### 3.1 Target selection

A set of single particle cryo-EM structures was chosen that would convey a representative sample of experimental reconstructions covering a wide range of nominal resolutions (d_min_) from 1.7-8.5 Å and symmetry conditions (1-24 symmetric copies). The test cases were restricted to EMDataBank (EMDB) (Lawson *et al*., 2016), entries for which half-maps had been deposited. Table 1 shows a summary of the selected test cases.

**Table 1.** Cryo-EM structures and models used for docking tests

| Target | code<sup>*</sup> | d<sub>min</sub> | copies<sup>†</sup> | model<sup>‡</sup> | fraction | model type |
| --- | --- | --- | --- | --- | --- | --- |
| GABA receptor | 7a5v_11657 | 1.7 | 5 | 4cofA (291-447)<br>(membrane domain) | 0.055 | crystal structure |
|  |  |  |  | AF model of megabody | 0.051 | AF prediction |
| Beta-galactosidase | 5a1a_2984 | 2.2 | 4 | 1jz7A | 0.25 | crystal structure |
|  |  |  |  | 5a1aA | 0.25 | reference |
|  |  |  |  | 5a1aA beta-barrel domain (626-726) | 0.025 | reference |
| Apoferitin | 5xb1_6714 | 3.0 | 24 | 5xb1A | 0.042 | reference |
|  |  |  |  | 2cei (5-159) | 0.042 | crystal structure |
| Respiratory complex | 7nyu_12654 | 3.8 | 1 | 3rkoAJKLMN<br>(membrane domain) | 0.42 | crystal structure |
|  |  |  |  | 3rkoN | 0.11 | crystal structure |
|  |  |  |  | 3rkoM | 0.11 | crystal structure |
|  |  |  |  | 3rkoL (1-546) | 0.11 | crystal structure |
|  |  |  |  | 7nyuM | 0.11 | reference |
| MutS | 7ai6_11792 | 6.9 | (2) | 6i5f (A12-A116) | 0.057 | crystal structure |
|  |  |  |  | 6i5f (A566-A799) | 0.13 | crystal structure |
| CFTRΔF508 mutant | 8ej1_28172 | 6.9 | 1 | AF (4- 263, 282-379, 844-871, 933-1170)<br>(membrane domain) | 0.52 | AF, no template |
|  |  |  |  | AF (264-281, 1204-1429) | 0.19 | AF, no template |
|  |  |  |  | AF (391-400, 440-633) | 0.17 | AF, no template |
| Get3, closed | EMD_25375 | 8.46 | (2) | 7spz<sup>¶</sup> | 0.5 | crystal structure |

<sup>\*</sup> Code for PDB plus EMDB pair, with the PDB identifier followed by the EMDB deposition number.
<sup>†</sup> Number of symmetry-related copies (or pseudo-symmetric copies if in parentheses).
<sup>‡</sup> Models from PDB entries are defined in terms of the PDB identifier, optionally followed by a chain identifier and/or a range of residue numbers. AF indicates AlphaFold model.
<sup>||</sup> Fraction of the entire ordered volume explained by one copy of the model.
<sup>¶</sup> Structure of Get3 in the open conformation.

### 3.2 Model selection

Models were selected to cover a variety of scenarios. Some models correspond to what could be called “reference” models, in the sense that they are the deposited models associated with the EMDB entry; these provide a reference docking with nearly zero rotation or translation. Others correspond to crystal structures of the same protein. Finally, we have tested some predicted models produced by AlphaFold (Jumper *et al*., 2021) (AF); such models will be used frequently, so understanding how they should be treated and how they will perform in our algorithm is essential. In all cases, we processed the predicted models with the *process_predicted_model* tool (Oeffner *et al*., 2022), which replaces the predicted values for the local distance difference test (Mariani *et al*., 2013), or pLDDT values, in the B-factor field of the coordinate file with appropriate B-factors to down-weight the less-confident parts of the model, as well as trimming off residues with a pLDDT value less than 70 (on a scale of 0-100).

To determine the effect of model completeness, as well as local map quality, we also tested the effect of using smaller pieces of the structural model (individual chains, domains or sub-domains). The models are also summarised in Table 1.

## 4. Implementation of algorithms

The algorithms have been implemented as a combination of Python scripts and C++ code, both making substantial use of the Computational Crystallography Toolbox, cctbx (Grosse-Kunstleve *et al*., 2002).

The framework for the docking search has been implemented in the Python program *em_placement*, which is part of the Voyager structural biology framework built on *phasertng* (McCoy *et al*., 2021). Associated tools required to evaluate the map eLLG, map information gain, fast phased translation search and cryo-EM likelihood target have been added to *phasertng*, which already contained tools to compute the rotation function eLLG (McCoy *et al*., 2017), fast searches and LLG rescoring for rotations (Storoni *et al*., 2004), and phased rigid-body refinement (Millán *et al*., 2021).

Note that the symmetry of the reconstruction is not yet used to aid model placement in the current version of the program.

The *em_placement* program is controlled using a set of keywords in the *phil* syntax used in Phenix (Liebschner *et al*., 2019). An example keyword script is given in Appendix A. Most keywords (map files, model file, composition of the reconstruction defined in terms of sequences of the components) will not usually be altered. The nominal resolution of the map is optional but recommended, and the author-defined value in the EMDB entry was used in all cases reported here. It would be appropriate to use either the FSC derived overall resolution or the highest local resolution in the map. Since the nominal resolution is used as the high-resolution limit for all the calculations, if the map actually contains valid higher resolution features than the user-entered value, some signal will be lost. If the user-entered value extends beyond the real resolution limit, CPU time is wasted but the search results should not be degraded unless the nominal resolution is very over-optimistic. The only parameter that might usefully be varied by the user is the equivalent RMS error that defines the expected model quality. For the test cases, a value of 0.8 Å was used for models obtained from experimental structures of the same protein, 1.0 Å for models predicted by AlphaFold, and 1.0 Å for the one experimental structure that differs somewhat in sequence, *i*.*e*. the model of apoferritin derived from a structure that contains the helix deleted in the target structure.

Note that the estimate of the equivalent RMS error is refined as part of the rigid-body refinement so, as long as the same solutions are found in the search, the final result is the same.

Data used for test calculations are all available through the EMDataBank (Lawson *et al*., 2016). Cryo-EM and crystallographic models are available from the worldwide Protein Data Bank (Berman *et al*., 2007), except for the AF models, which were computed using the community ColabFold version (Mirdita *et al*., 2022) of AlphaFold (Jumper *et al*., 2021).

## 5. Results

### 5.1 Docking results

The results of the docking trials are summarized in Table 2. The majority of the searches succeeded, and many of these required only a single search over the entire reconstruction. The time required for the searches ranged from half a minute to about 32 minutes, averaging about 12 minutes over the set of test cases. When multiple spherical sub-volumes were searched, the number varied from 4 to 214. None of the test cases triggered the fall-back of carrying out a brute-force six-dimensional search.

**Table 2.**
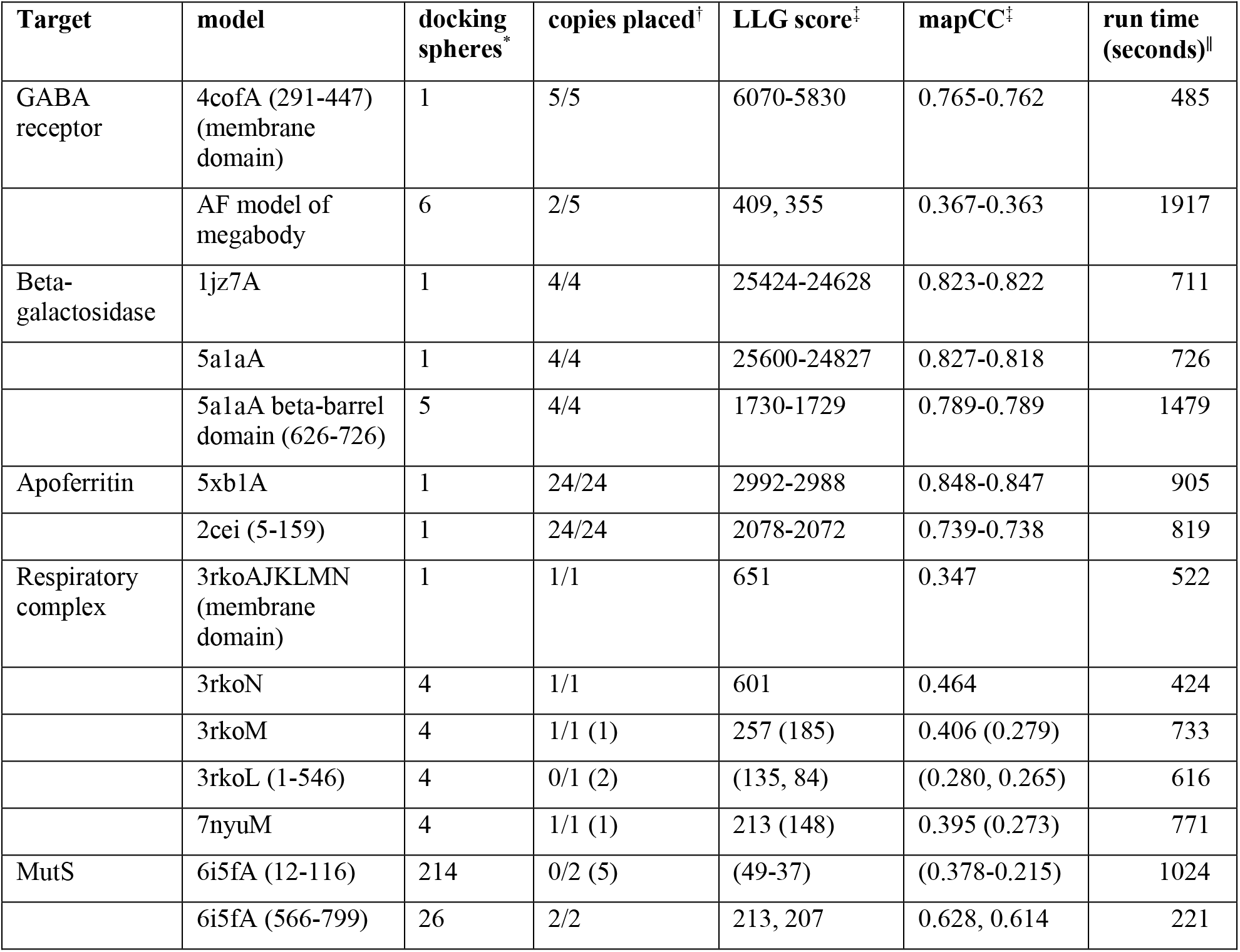
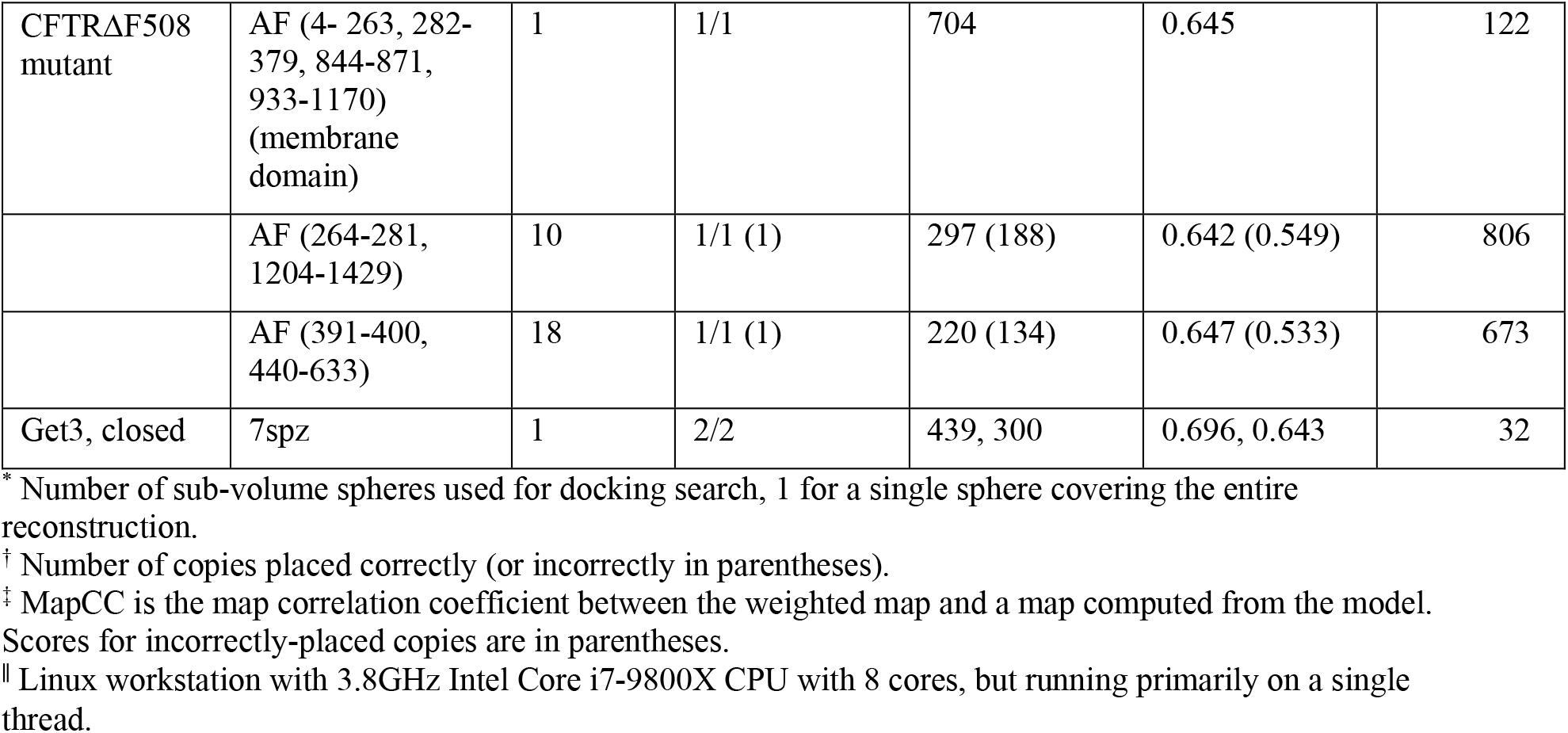
Results of docking trials

| Target | model | docking spheres* | copies placed† | LLG score‡ | mapCC‡ | run time (seconds)¶ |
| --- | --- | --- | --- | --- | --- | --- |
| GABA receptor | 4cofA (291-447) (membrane domain) | 1 | 5/5 | 6070-5830 | 0.765-0.762 | 485 |
|  | AF model of megabody | 6 | 2/5 | 409, 355 | 0.367-0.363 | 1917 |
| Beta-galactosidase | 1jz7A | 1 | 4/4 | 25424-24628 | 0.823-0.822 | 711 |
|  | 5a1aA | 1 | 4/4 | 25600-24827 | 0.827-0.818 | 726 |
|  | 5a1aA beta-barrel domain (626-726) | 5 | 4/4 | 1730-1729 | 0.789-0.789 | 1479 |
| Apo ferritin | 5xb1A | 1 | 24/24 | 2992-2988 | 0.848-0.847 | 905 |
|  | 2cei (5-159) | 1 | 24/24 | 2078-2072 | 0.739-0.738 | 819 |
| Respiratory complex | 3rkoAJKLMN (membrane domain) | 1 | 1/1 | 651 | 0.347 | 522 |
|  | 3rkoN | 4 | 1/1 | 601 | 0.464 | 424 |
|  | 3rkoM | 4 | 1/1 (1) | 257 (185) | 0.406 (0.279) | 733 |
|  | 3rkoL (1-546) | 4 | 0/1 (2) | (135, 84) | (0.280, 0.265) | 616 |
|  | 7nyuM | 4 | 1/1 (1) | 213 (148) | 0.395 (0.273) | 771 |
| MutS | 6i5fA (12-116) | 214 | 0/2 (5) | (49-37) | (0.378-0.215) | 1024 |
|  | 6i5fA (566-799) | 26 | 2/2 | 213, 207 | 0.628, 0.614 | 221 |
| CFTRΔF508 mutant | AF (4- 263, 282-379, 844-871, 933-1170) (membrane domain) | 1 | 1/1 | 704 | 0.645 | 122 |
|  | AF (264-281, 1204-1429) | 10 | 1/1 (1) | 297 (188) | 0.642 (0.549) | 806 |
|  | AF (391-400, 440-633) | 18 | 1/1 (1) | 220 (134) | 0.647 (0.533) | 673 |
| Get3, closed | 7spz | 1 | 2/2 | 439, 300 | 0.696, 0.643 | 32 |

\* Number of sub-volume spheres used for docking search, 1 for a single sphere covering the entire reconstruction.
† Number of copies placed correctly (or incorrectly in parentheses).
‡ MapCC is the map correlation coefficient between the weighted map and a map computed from the model. Scores for incorrectly-placed copies are in parentheses.
§ Linux workstation with 3.8GHz Intel Core i7-9800X CPU with 8 cores, but running primarily on a single thread.

#### 5.1.1 GABA receptor

The highest-resolution (1.7 Å) cryo-EM structure in our test set is that of the human γ-aminobutyric acid receptor bound to a megabody, PDB entry 7a5v, EMDB entry 11657 (Nakane *et al*., 2020).

To provide a reasonable challenge at such high resolution, only small models were tested, each comprising about 1/20 of the full pentamer or 1/4 of a single copy. The membrane domain is well-ordered and is easy to place when using the membrane component of a single subunit of a crystal structure, PDB entry 4cof (Miller & Aricescu, 2014), as a model. However, an AlphaFold model of the bound megabody is more difficult to place, as the associated density is the least well-ordered in the map. Only 2 of the 5 copies were placed successfully, in spite of the 5-fold symmetry of the reconstruction. If the sub-volume spheres were placed in a way that obeyed the 5-fold symmetry, the same results would be obtained for each copy. The sensitivity of the search to the boundaries of the sub-volumes is an indicator that this is a marginal model for searching in this map. In principle, the missing copies could be generated by application of the 5-fold symmetry.

#### 5.1.2 Beta-galactosidase

Beta-galactosidase is commonly used as a test object for cryo-EM methodology, as it is well-behaved and possesses D2 tetrameric symmetry. We chose a medium-resolution (2.2 Å) representative: PDB entry 5a1a, EMDB entry 2984 (Bartesaghi *et al*., 2015).

Docking a full chain, either from the associated PDB entry or from a crystal structure, PDB entry 1jz7 (Juers *et al*., 2001), is straightforward to achieve by searching over the full map.

On the other hand, docking just the beta-barrel domain of one subunit is substantially more challenging, and the map is divided into 5 sub-volumes. All 4 copies were found successfully, although this success is fragile. A parallel run under MacOS found only 3 copies using the same computer code, presumably because of the effects of minor numerical differences. Again, the missing copy in that case could have been recovered by exploiting the symmetry of the map.

#### 5.1.3 Apoferritin

Because of its stability and high octahedral (432) symmetry, apoferritin is another very common test object for cryo-EM. We chose a relatively low-resolution (3.0 Å) representative: PDB entry 5xb1, EMDB entry 6714, a deletion mutant of the E-helix (Ahn *et al*., 2018).

Searching with a single chain from the reference structure finds all 24 copies with strong signal in a search over the full volume; the default to keep a maximum of 5 potential solutions was overridden for this case. As a more challenging test, we based a search model on a single chain from PDB entry 2cei, the crystal structure of a full-length version of apoferritin, removing the E-helix from the search model. Again, all 24 copies were found with strong (though slightly lower) signal. Note that much of the computing time in these two tests is expended on evaluating the map correlations for the 24 solutions.

#### 5.1.4 *E. coli* respiratory complex I

The largest series of trials was carried out with the reconstruction for conformation 2 of the *E. coli* respiratory complex I: PDB entry 7nyu, EMDB entry 12654 (Kolata & Efremov, 2021). This reconstruction presents a variety of challenges, as the overall resolution (3.8 Å) is already relatively low but also varies substantially over the different subunits. Parts of the membrane domain are particularly poorly resolved; the local resolution of chain L is estimated by the authors as being in the range of 9-11 Å. An additional challenge comes from the fact that three of the membrane domain components (chains L, M and N) have related sequences and structures, with pairwise sequence identities of 25-26%. As a result, it is possible to place a model into the density for a related subunit, yielding a non-random LLG score above 60.

Models were taken either from the reference structure or from the crystallographic structure of the membrane domain, PDB entry 3rko (Efremov & Sazanov, 2011). Searching for the entire membrane domain gives a clear solution using the full reconstruction. In searches for individual chains, such as the three related membrane domain components, the reconstruction is automatically divided into sub-volumes. For the best-ordered of the three related subunits, chain N, an unambiguous solution is found. Chain M is more poorly-ordered, and two potential solutions are found. The solution with higher scores is correctly placed, while the second solution superimposes the chain M model on the better-ordered density of chain N. Chain L is the least well-ordered, and the search places the model on either the density for chain N or chain M, but not on the correct density that corresponds to chain L. Fig. 1a illustrates one of the most difficult successful results, showing the docked model of chain M.

**Figure 1.**
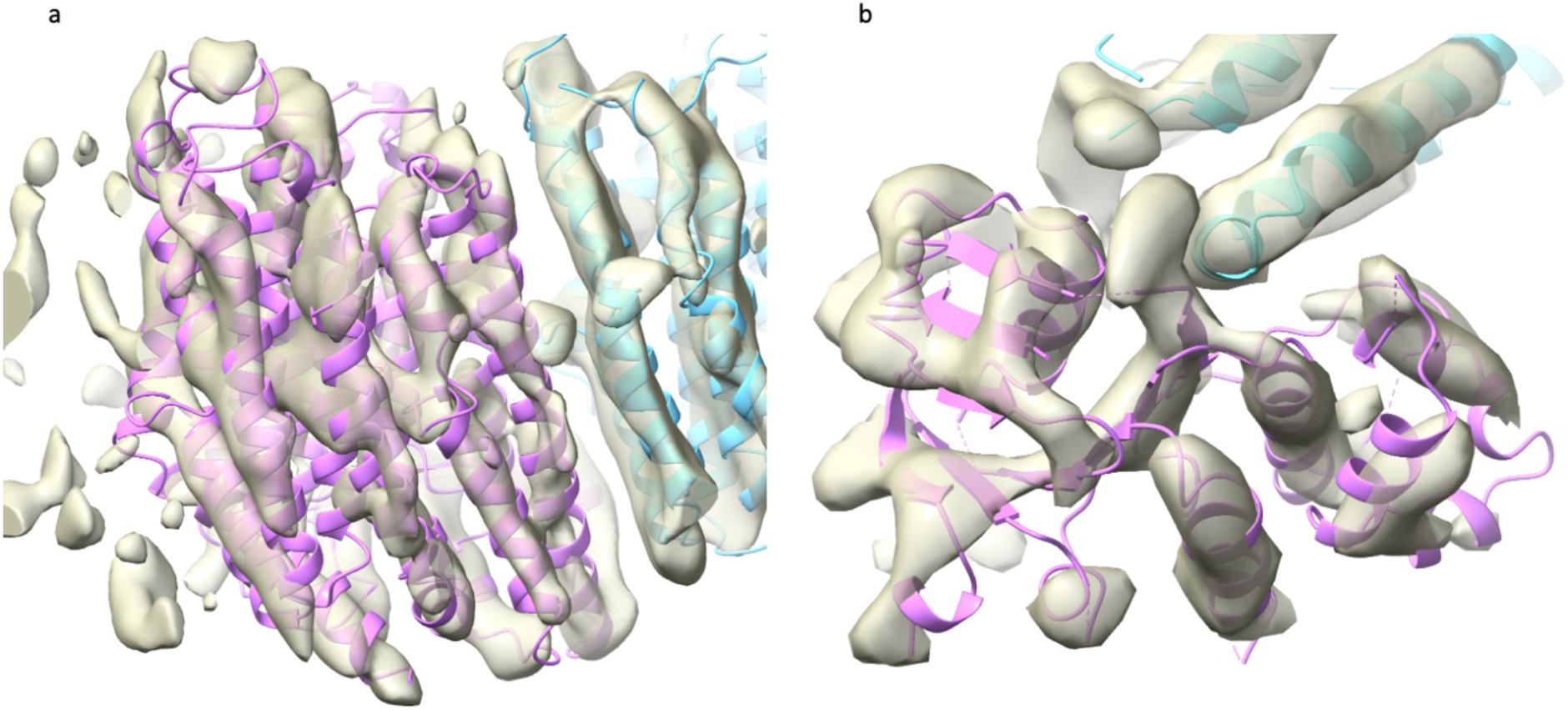
Docked models in maps for challenging cases. Both maps are computed using the Fourier coefficients 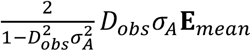 arising from the analysis of the local map volumes, and the images were made with ChimeraX (Goddard *et al*., 2018). a) Chain M (magenta) of PDB entry 3rko, docked into the region of the map corresponding to chain M of PDB entry 7nyu (associated with EMDB entry 12654). Chain N is shown in light blue. b) The AlphaFold model of the smallest domain of the ΔF508 mutant of CFTR (magenta), docked into the corresponding region of the map derived from EMDB entry 28172. The membrane domain is shown in light blue.

#### 5.1.5 DNA mismatch repair protein, MutS

For one representative of a low-resolution (6.9 Å) reconstruction, we chose the *E. coli* DNA mismatch repair protein, MutS, in its mismatch-bound state: PDB entry 7ai6, EMDB entry 11792. In this bound state, the protein is a pseudosymmetric dimer, so there are two independent copies to find.

To test a workflow in which individual domains are docked, in order to approximate a conformational change, we used as models the N- and C-terminal domains of one chain of MutS in the DNA-free conformation, from PDB entry 6i5f (Bhairosing-Kok *et al*., 2019). For such small fractions of the full structure at such low resolution, the signal in the rotation search would be extremely low, so the sub-volume determination algorithm chose to carry out the searches with multiple sub-volumes of the maps. For the (larger) C-terminal domain, 26 spherical sub-volumes were chosen. Although this is a relatively large number, each calculation is fast with low-resolution data, and an unambiguous docking of both copies was achieved in less than 4 minutes. For the (smaller) N-terminal domain, 214 sub-volumes were chosen. The search in this case took significantly longer, at about 17 minutes, and it failed to find the correct placements. None of the LLG values exceeded 50.

#### 5.1.6 Cystic fibrosis transmembrane regulator, ΔF508 mutant

The ΔF508 mutant of the cystic fibrosis transmembrane regulator (CFTR), with bound folding modulators, was chosen as a second low-resolution (6.9 Å) reconstruction: PDB entry 8ej1, EMDB entry 28172 (Fiedorczuk & Chen, 2022).

Rather than testing other experimental structures of the same protein, we chose to make AlphaFold (Jumper *et al*., 2021) models in the ColabFold environment (Mirdita *et al*., 2022). Although structures of the CFTR would have been present in the training data for AlphaFold, their influence was reduced by turning off the option to include explicit templates of related structure in the structure prediction process. As for the MutS case, the difficulty of the docking calculations was increased by extracting models of individual domains from the full predicted structure. As expected, it was more difficult to place smaller models. The membrane domain, the largest with 585 residues in the processed model, was placed easily (LLG = 704) in a search over the entire reconstruction. A mid-sized domain, comprising 214 residues, gave two potential solutions with LLG values of 297 and 188 but searching over 10 sub-volumes and taking nearly seven times as long. The first potential solution was placed correctly, but the second was superimposed on the smallest domain, which has a similar fold and a sequence identity of 27% over 168 matched residues (of 187 in the smaller of the two domains). Similarly, a search for the smallest domain gave two potential solutions, with LLG scores of 220 for the correct solution (Fig. 1b) and 134 for a superposition on the mid-sized domain.

#### 5.1.7 Get3, closed conformation

The lowest-resolution (8.46 Å) map in the test set is of the closed conformation of the ER targeting factor Get3: EMDB entry 25375 (Fry *et al*., 2022). The authors did not deposit coordinates in the PDB for this reconstruction, presumably because it had the lowest resolution of a series of maps. Therefore, it makes a good example for the circumstance in which a structural biologist would like to examine a published map in the context of a docked model from another structure.

We chose the crystal structure the same authors determined for the open conformation, from PDB entry 7spz (Fry *et al*., 2022), as the model. The reconstruction is pseudosymmetric, so there are two independent copies to find. Both of them can be found in a straightforward search over the full reconstruction that takes only about half a minute.

### 5.2 Checking the *eLLG*_*rot*_-guided sub-volume criterion

In the global search, the *eLLG*_*rot*_ criterion suggested that a single search sphere covering the entire ordered volume of the reconstruction would give sufficient rotation function signal for 8 of the 18 test cases. Validating this, all 8 of these searches succeeded (Table 2). However, if the *eLLG*_*rot*_ criterion were too pessimistic about the ability to find the model in the whole map, the 8 searches that found at least one copy when searching over sub-volumes might have succeeded with a global search. To test this, we used a manual override in the *em_placement* program to force a search over the single sphere covering the entire ordered volume. Because the two models for chain M of the *E. coli* respiratory complex I are very similar, we only tested chain M from the crystal structure in PDB entry 3rko. The results are given in Table 3.

**Table 3.**
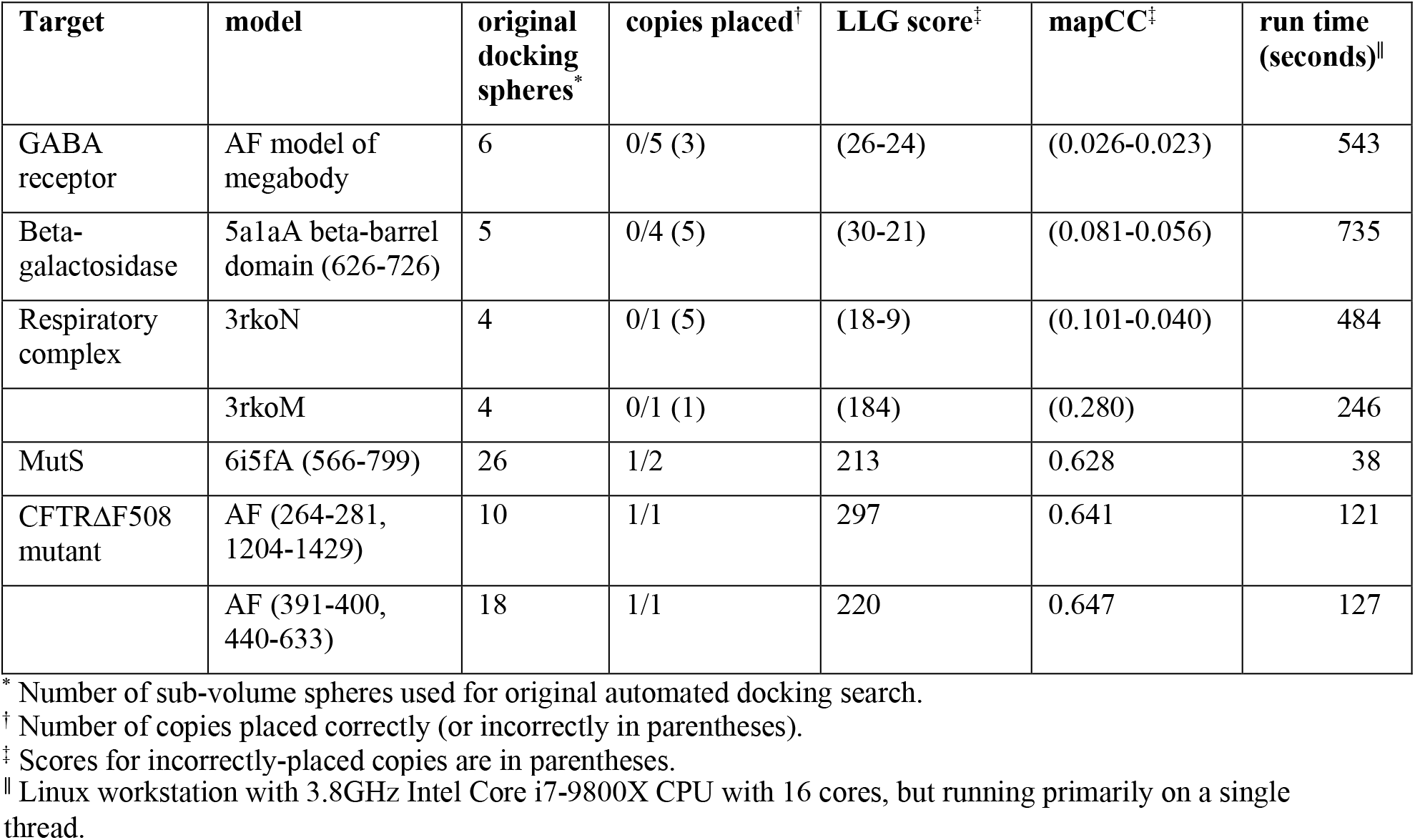
Results of trials searching globally over a single sphere

The results support the *eLLG*_*rot*_ criterion as an effective guide to search strategy. No correct solution is found for four of the seven test cases, and only one of two solutions is found when searching for the C-terminal domain of MutS. The only cases where the criterion was clearly too pessimistic about the ability to find the model in the whole map are the searches for the mid-sized and smallest domains of the AlphaFold model for the ΔF508 mutant of CFTR. Here the correct solutions are found in about two minutes each in the whole map, whereas the global sub-volume searches took 11-13 minutes (Table 2). However, the forced global search failed to find the non-random solutions mentioned in section 5.1.4 in which the two homologous domains were placed in positions belonging to each other..

### 5.3 Tests of brute-force six-dimensional searches

The two cases where the global search failed, as well as the MutS case in which 26 sub-volumes were explored, provided tests of the brute-force 6D fall-back algorithm. These were carried out to examine whether the global 6D search could succeed for cases where rotation searches for the smallest practical sub-volume would have insufficient signal, and also how it compares in efficiency to searching over a large number of sub-volumes.

#### 5.3.1 Chain L of the *E. coli* respiratory complex I

The brute-force 6D search fails to find the correct position of chain L, but does reproduce the results of the automated search using multiple sub-volumes as the model for chain L is superimposed on the map regions for chains M and N. The run time is dramatically longer at approximately 11 hours, compared to about 10 minutes for the automated search with multiple sub-volumes.

#### 5.3.2 N-terminal domain of MutS

The brute-force search is more successful than the adaptive search over 214 sub-volumes, as one of the two copies of this domain is found with an LLG of 83 and a map correlation of 0.536. Although the next best potential solution has an LLG of 60, none of the other potential solutions are correct. The partial success comes at the cost of about 44 minutes running time. This is the only test case in which triggering the fall-back to a brute-force 6D search would have been justified.

#### 5.3.3 C-terminal domain of MutS

Both copies of the C-terminal domain of PDB entry 6i5f are found in the brute-force 6D search, with the same scores. However, the search using 26 sub-volume spheres is dramatically more efficient, taking about 4 minutes compared to 139 minutes for the brute-force 6D search.

## 6. Discussion and conclusions

The strength of likelihood as a criterion is supported by the success of our new likelihood-based approach to docking models in a series of progressively more challenging cryo-EM maps. Since the successful application of likelihood to a problem requires a good model of the sources of error and their propagation, these results also support our approach to defining and calibrating likelihood targets for cryo-EM data in the accompanying paper (Read *et al*., 2023).

The outcomes of different search strategies can be predicted by an analysis of the expected log-likelihood-gain (eLLG) score for both the rotation and translation search components of the docking algorithm. The rotation function eLLG can be used to predict how large a volume of the map can be explored in one rotation search, allowing automated decisions about the subdivision of the full map into spherical sub-volumes. The choices made by this criterion have been validated by comparing the success of searches over the full map with those carried out over the suggested sub-volumes.

Docking models into the most poorly-ordered part of a map is difficult, partly because of the reduced signal to noise but also because the assessment of global map quality can mislead the algorithm determining search strategy into choices that provide insufficient signal in the worst regions of the map. This could potentially be mitigated by adapting the strategic choices to local levels of signal to noise in the reconstruction.

Plans for future enhancements include accounting for symmetry in the search space, which will be significantly more efficient in the case of high-symmetry reconstructions such as the ones for apoferritin. Calculations could be made faster by using parallel processing, particularly for searches over multiple sub-volumes. Searches for multiple components will be implemented, which requires accounting for the contribution of previously-placed components in the fit to the experimental data, as well as avoiding clashes between components.

## Appendix A.

**Example script for *em_placement***

The following script defines the search parameters for the *em_placement* script used to run the first test case, docking the membrane component of a model of the GABA receptor derived from PDB entry 4cof into the cryo-EM reconstruction deposited as EMDB entry 11657.

~~~
voyager
{
map_model
{
half_map = emd_11657_half_map_1.map
half_map = emd_11657_half_map_2.map
best_resolution = 1.7 point_group_symmetry = C5
sequence_composition = 7a5v.fa
}
biological_unit {
molecule
{
molecule_name = 4cofA_membrane
map_or_model_file = 4cofA_membrane.pdb
starting_model_vrms = 0.8
}
}
}
~~~

Using a recent *Phenix* version (the dev-4887 development release or newer), this script can be run with the command:

phenix.voyager.em_placement docking_script.phil.

Most parameters specified in the script have been named in a way intended to convey the purpose of that parameter. The point_group_symmetry feature is only used at the moment to optionally generate a full assembly from a single copy. The sequence_composition parameter specifies the name of a file containing the sequences of all the components in the reconstruction.

## Acknowledgements

We thank Cathy Lawson for implementing a method to allow searches at the EMDataResource for EMDB entries providing half-maps.

